# Clinical positioning of the IAP antagonist tolinapant (ASTX660) in colorectal cancer

**DOI:** 10.1101/2020.11.20.391680

**Authors:** Nyree Crawford, Katie Stott, Tamas Sessler, Christopher McCann, William McDaid, Cheryl Latimer, Jennifer Fox, Joanne M Munck, Tomoko Smyth, Alpesh Shah, Vanessa Martins, Mark Lawler, Philip Dunne, Emma Kerr, Simon S McDade, Vicky Coyle, Daniel B Longley

## Abstract

Cancer cells frequently express elevated levels of Inhibitor of Apoptosis Proteins (IAPs): cIAP1, cIAP2 and XIAP. Elevated expression of cIAP1 and cIAP2 (but not XIAP) significantly correlated with poor prognosis in microsatellite stable (MSS) stage-III colorectal cancer (CRC) patients treated with adjuvant chemotherapy, suggesting their involvement in promoting resistance. Preclinical analysis of the IAP inhibitor tolinapant in CRC cell lines demonstrated robust on-target effects and caspase-8-dependent apoptosis that was inhibited by the caspase-8 paralogs FLIP and, unexpectedly, caspase-10. Importantly, tolinipant-induced apoptosis was augmented by standard-of-care chemotherapy (FOLFOX) in CRC disease models, due (at least in part) to FOLFOX-induced downregulation of Class-I histone deacetylases, leading to acetylation of the FLIP-binding partner Ku70 and downregulation of FLIP. Moreover, this effect could be phenocopied using a Class-I HDAC inhibitor. Further analyses revealed that caspase-8-knockout RIPK3-positive CRC models were sensitive to tolinostat-induced necroptosis, an effect that could be exploited with the FDA-approved caspase inhibitor emricasan. Our study provides evidence for immediate clinical exploration of tolinapant in combination with FOLFOX chemotherapy in poor prognosis MSS CRC with elevated cIAP1/2 expression.

## Introduction

Colorectal cancer (CRC) is the second leading cause of cancer-related deaths in the Western world. In stage IV metastatic disease, 5-year and 10-year overall survival rates are <6% and <1% respectively [1]. Fluoropyrimidine-based chemotherapy is the mainstay of standard-of-care (SoC) treatment for CRC in both early and advanced disease. However in early stage disease, the benefit of fluoropyrimidine monotherapy is limited, with overall survival gains of less than 5% [2] and 10-15% [3] reported for Stage II and III disease respectively, with a small additional survival gain for use in combination with oxaliplatin [4]. Improved approaches are therefore urgently needed not only for the treatment of advanced CRC, but also in the adjuvant disease setting for those patients who are destined to relapse but whose disease is resistant to SoC chemotherapy.

Cancer cells are frequently ‘addicted’ to a small number of anti-apoptotic proteins that are necessary for their survival. These include the Inhibitor of Apoptosis Proteins (IAPs), cIAP1, cIAP2 and XIAP [5]. XIAP directly binds to, and inhibits the activation of, caspases-3, −7 and −9 [6]. cIAP1 and cIAP2 act as E3 ubiquitin ligases at Complex I which is formed following binding of the pro-inflammatory cytokine, TNFα, to TNF receptor 1 (TNFR1). cIAP1 and cIAP2 polyubiquitinate RIPK1 resulting in the activation of the canonical nuclear factor-κB (NFκB) signalling pathway [7], which in turn results in the transcription of anti-apoptotic proteins, including cIAP1, cIAP2 and inflammatory cytokines (including TNFα itself) in a pro-survival/inflammatory feedforward loop [8, 9]. Complex I dissociates following deubiquitination of RIPK1 to form the death-inducing platform, Complex IIa composed of FADD, RIPK1, TRADD and procaspase-8 [10]. In the absence of cIAPs, RIPK1 is not polyubiquitinated, and the cell death inducing platform complex IIb (containing of FADD, RIPK1 and caspase-8) is formed, which can also activate cell death. Hence targeting IAPs have potential to both activate cell death and inhibit anti-apoptotic and pro-inflammatory signalling.

Multiple peptidomimetic IAP antagonists based on the endogenous inhibitor SMAC (second mitochondrial activator of caspases) have been explored in the clinical setting. These compounds have inherent selectivity for cIAP1 over XIAP. Tolinapant (ASTX660) (Astex Pharmaceuticals) is a novel, non-peptidomimetic IAP antagonist with balanced nanomolar potency against cIAP1/2 and XIAP that was discovered utilising fragment-based drug design [11]. The first-in-human phase 1 trial of tolinapant indicated a manageable safety profile and had evidence of clinical activity; phase 2 studies are ongoing (NCT02503423) [12].

In this study, we assessed the activity of tolinapant in pre-clinical models of CRC as a single agent and in combination with both SoC chemotherapeutics and novel therapies rationally selected based on mechanistic insights.

## Results

### Elevated cIAP1 and cIAP2 expression correlates with worse prognosis of MSS stage-III CRC patients treated with adjuvant chemotherapy

Analysis of CRC patients [13] revealed that elevated expression of *cIAP1/BIRC2* (HR=0.47; p=0.05) and *cIAP2/BIRC3* (HR=0.34; p=0.009)(which are normally distributed (**Figure S1A**)), but not *XIAP/XIAP* correlated with worse overall survival (OS) in MSS stage-III patients treated with chemotherapy post-surgery (adjuvant chemotherapy) (**Figure 1A** and **Figure S1B**), whereas no significant association between IAP expression and patient survival in stage-II disease, nor in stage-III patients treated by surgery alone (*data not shown*). While cIAP-1 and cIAP-2 expression were significantly correlated (Pearson’s r = 0.47, p = 1.3e(−8)), not all cIAP1-high tumors were also cIAP2-high (**Figure 1C and S1C**). Single sample gene set enrichment analyses (ssGSEA) comparing cIAP-1-or cIAP-2-high and -low groups identified significant enrichments for several key inflammatory signalling pathways in both the cIAP1- and cIAP2-high groups, including *“TNFα* signalling via *NFκB*” and other inflammatory pathways (**Figure 1B/C**). Consistent with this, the tumours from the cIAP1- and cIAP2-high groups were enriched for the presence of several key immune cell types, with significant enrichment of CD4+ memory T cells, TH2 cells and M1 polarised macrophages in both groups (**Figure 1D/E**). In addition to apoptosis signalling, there was also a significant enrichment for KRAS signalling in both the cIAP1- and cIAP2-high groups (**Figure 1B**). IAP expression was not higher in *KRAS* mutant tumours (and was not correlated with *TP53* or *BRAF* mutations; **Figure S1C/D**), indicating that the correlations between cIAP1/2 expression and survival (**Figure 1A**) are not due to enrichment for *KRAS* mutant tumours, which have a worse, but not significantly worse prognosis in this MSS stage-III cohort (**Figure S1E**).

**Figure 1:**
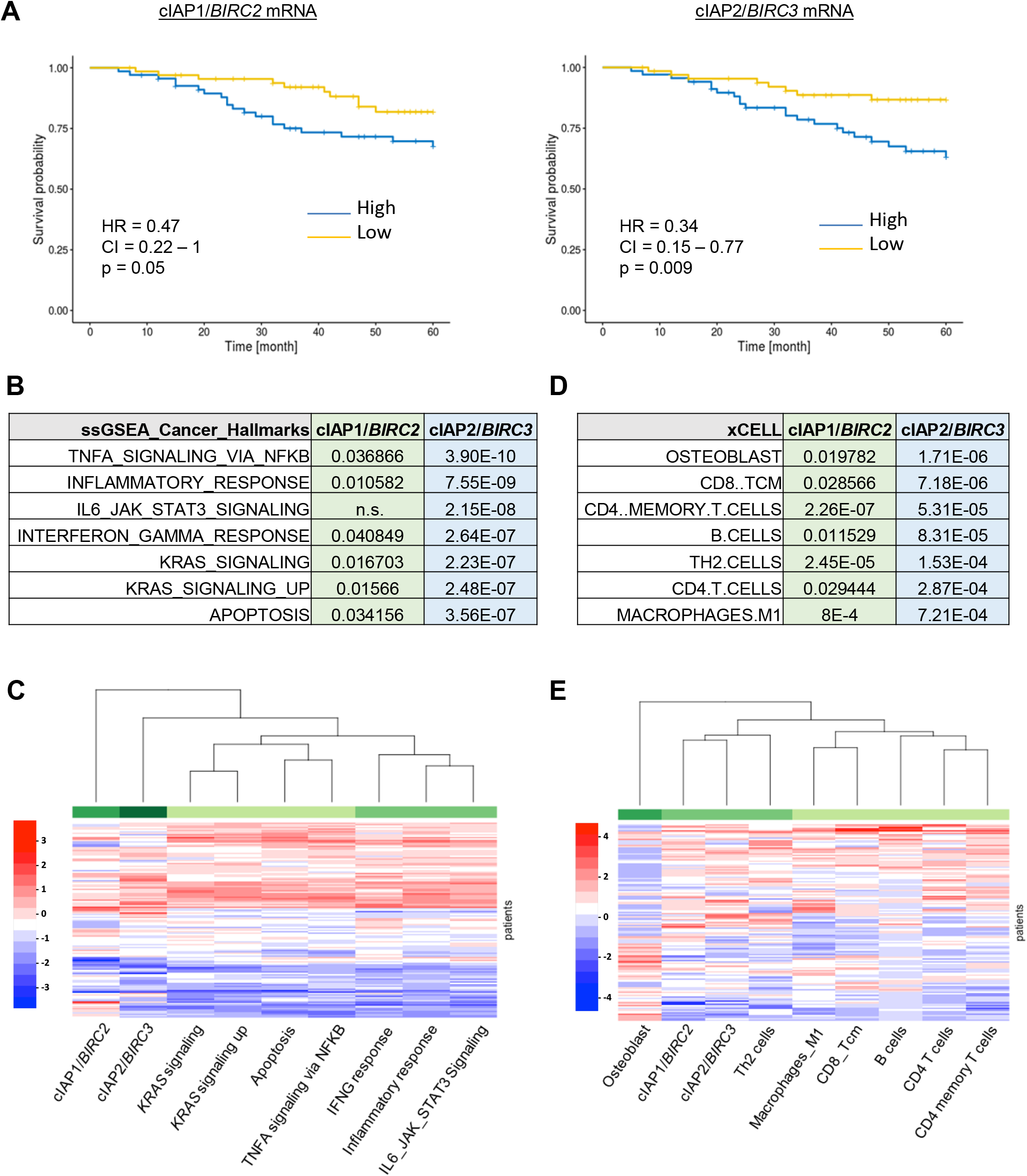
Analysis of MSS, Stage III patients who received adjuvant fluorouracil-based treatment extracted from GSE39582 cohort. **A)** Kaplan-Meier plots of 5-year overall survival. Graphs comparing the difference in survival probability based on BIRC2 and BIRC3 expression levels alone. Results are shown with the log-rank test, the hazard ratio (HR) with its 95% confidential interval and the Wald test of significance. **B-E)** Difference of pathway activation by **B)** ssGSEA and **D)** cell type profiling by xCELL between cIAP1 or cIAP2 high and low expressing patients was assessed by one-way anova model. Numbers shown are the calculated p values. Heatmap were drawn showing the Z-scores of the significantly different **C)** ssGSEA pathways and **E)** immune or stroma infiltration by xCELL.

### Sensitivity of CRC cell lines to tolinapant in vitro and in vivo

The analysis of the clinical cohort suggested potential for cIAP1 and cIAP2 as potential therapeutic targets for improving the efficacy of standard-of-care chemotherapy in MSS stage-III disease. We therefore next assessed the cell death inducing activity of the novel IAP antagonist tolinapant by screening a panel of 36 CRC cell lines alone or in the presence of low concentrations of TNFα (**Figure 2A**); to reflect the enrichment for TNFα signalling in the cIAP1/2-high groups (**Figure 1B/C**) and the presence of immune cells such as M1 macrophages (**Figure 1D/E**), which are rich sources of TNFα [14]. In the absence of TNFα, tolinpant had modest growth inhibitory effects across the panel, whereas in the presence of TNFα, a high number of cell lines exhibited significant sensitivity to tolinpant (**Figure 2A**). These effects were further assessed in a subset of CRC cell line models with a range of sensitivities (**Figure S2A**), each of which expressed all the components of the TNFR1 Complex-II interactome and XIAP at the protein level (albeit to varying levels; **Figure 2B**) as well as cell surface TNFR1 (**Figure 2C**). On-target activity of tolinapant as evidenced by degradation of cIAP1 (the pharmacodynamic (PD) biomarker for IAP antagonists), was confirmed at nM potency in all 5 models (**Figure 2D** and **Figure S2B**); whereas cIAP2 was also downregulated (albeit to a lesser extent than cIAP1) by tolinapant in 3/5 models. XIAP was unaffected, consistent with the expected effects of IAP antagonists which disrupt XIAP’s protein-protein interactions with caspases-3, −7 and −9, but do not typically cause XIAP degradation [15]. Downregulation of cIAP1 in response to tolinapant, which is a result of triggering its auto-ubiquitination and degradation via the proteasome [16], was found to be rapid and sustained (**Figure S2C**). Loss of IAPs has been widely reported to result in the formation of the cell death platform, Complex-II [17]. In the presence of TNFα, tolinapant indeed promoted the formation of complex-II as evidenced by the interaction of caspase-8 with RIPK1; this was also detected in response to tolinapant alone in 3/5 models HCT116, HT29 and LoVo cells albeit to a much lower level than in response to tolinapant/TNFα (**Figure 2E** and **Figure S2D**).

**Figure 2:**
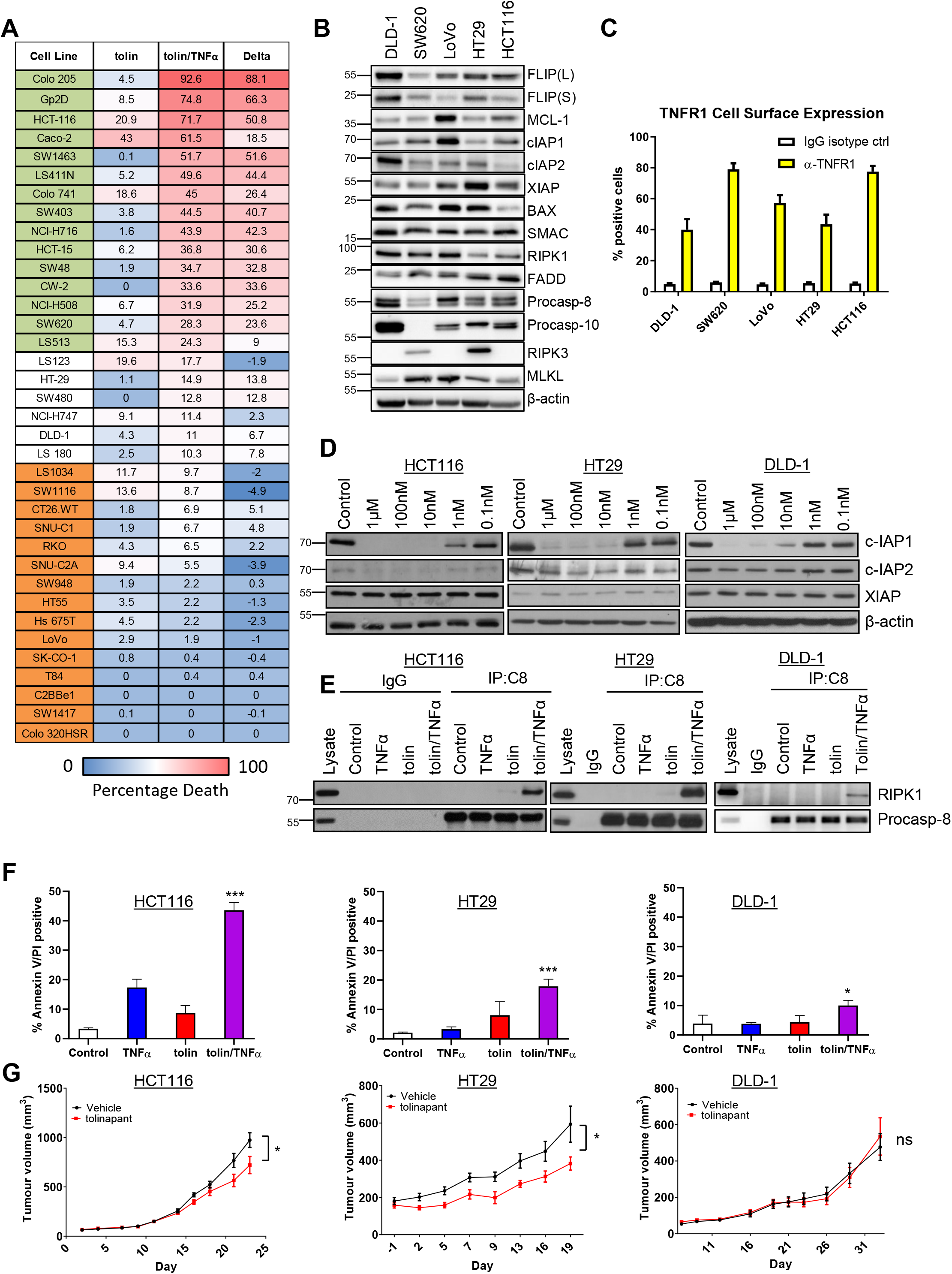
**A)** Cell viability assay in 36 CRC cell line panel following 72h treatment with tolinapant(tolin) (10 point assay 0.0005 – 10μM) alone and in combination with TNFα (1ng/ml), represented as relative activity area (%), calculated as (experimental AUC)/max AUC possible). Cell lines categorized as sensitive to tolinapant/TNFα are highlighted in green, intermediate in white and resistant in orange. **B)** Western blot analysis of basal expression of FLIP(L), FLIP(S), Mcl-1, cIAP1, cIAP2, XIAP, BAX, SMAC, RIPK1, FADD, Procaspase-8, Procaspase-10, RIPK3 and MLKL and β-actin in a panel of CRC cell lines. **C)** Cell surface expression of TNFR1 in DLD1, SW620, LoVo, HT29 and HCT116 cell lines, represented as % of TNFR1 positive cells compared to an IgG isotype control. **D)** Western blot analysis of cIAP1, cIAP2, XIAP and β-actin in HCT116, HT29 and DLD-1 cells following treatment with 1μM, 100nM, 10nM, 1nM and 0.1nM tolinapant for 24h. **E)** Western blot analysis of RIPK1 and procaspase-8 following immunoprecipitation of caspase-8 from HCT116, HT29 and DLD-1 cells treated with 1ng/ml TNFα, 1μM tolinapant and a combination of TNFα/ tolinapant for 3h in the presence of 20μM z-VAD-fmk. An IgG isotype control was used as a control. **F)** AnnexinV/PI analysis in HCT116, HT29 and DLD-1 cells following treatment with 1ng/ml TNFα, 1μM tolinapant or a combination of TNFα/tolinapant for 24h. **G)** Tumor volume (mm^3^) in HCT116, HT29 and DLD-1 xenograft models treated with vehicle and 16mg/kg tolinapant. Data shows mean tumor volume per treatment group per timepoint ± sem (HCT116; Vehicle n=10, tolinapant n=10, HT29; Vehicle n=7, tolinapant n=7, SW620; Vehicle n=10, tolinapant n=12). Results were compared using a two-tailed Students t-test, p<0.05 *, p<0.01 ** and p<0.001 ***.

In agreement with the cell line screen, significant cell death measured by Annexin-V/PI flow cytometry, was observed in response to tolinapant in the presence of TNFα in the HCT116 model (**Figure 2F**). Similar results were observed in the HT29 model, albeit the extent of cell death at 24h was less; however, lower levels of cell death were observed in the DLD-1 model despite complex-II formation (**Figure 2E/F**). In the *in vivo* setting, we hypothesised that murine TNFα (as well as xenograft-derived human TNFα) could potentially drive complex-II formation as murine TNFα can bind the human TNFR1 [18]. Moreover, IAP antagonists have been reported to promote production of TNFα by monocytic cells [19]. Significant albeit limited retardations of tumor growth were observed in HCT116 and HT29 xenografts but not DLD-1, results which were in line with the relative *in vitro* sensitivity of these models (**Figure 2G**).

### Standard-of-care chemotherapy enhances sensitivity to tolinapant

Having established the effects of tolinapant/TNFα combinations, we next evaluated whether the addition of SoC chemotherapy FOLFOX (5-Fluorouracil plus Oxaliplatin) could increase sensitivity to tolinapant. Statistically significant enhancement of FOLFOX-induced apoptosis was observed in 3/5 models (HCT116, SW620 and HT29) co-treated and tolinapant/TNFα (**Figure 3A/B**). Indeed, tolinapant alone also significantly enhanced FOLFOX-induced apoptosis in the HCT116 and SW620 models, albeit to a lesser extent than in the tolinapant/TNFα combination group (**Figure 3A/B**). In contrast, the DLD1 model was relatively resistant to the combination (**Figure S3A**), whereas interestingly, LoVo cells, which were relatively insensitive to the tolinapant/TNFα combination alone, exhibited significantly increased chemotherapy-induced apoptosis when co-treated with tolinapant alone as well as tolinapant/TNFα (**Figure S3A/B**). These results were reflected by qualitative assessment of apoptosis by PARP cleavage (**Figure 3B/S3B**).

**Figure 3:**
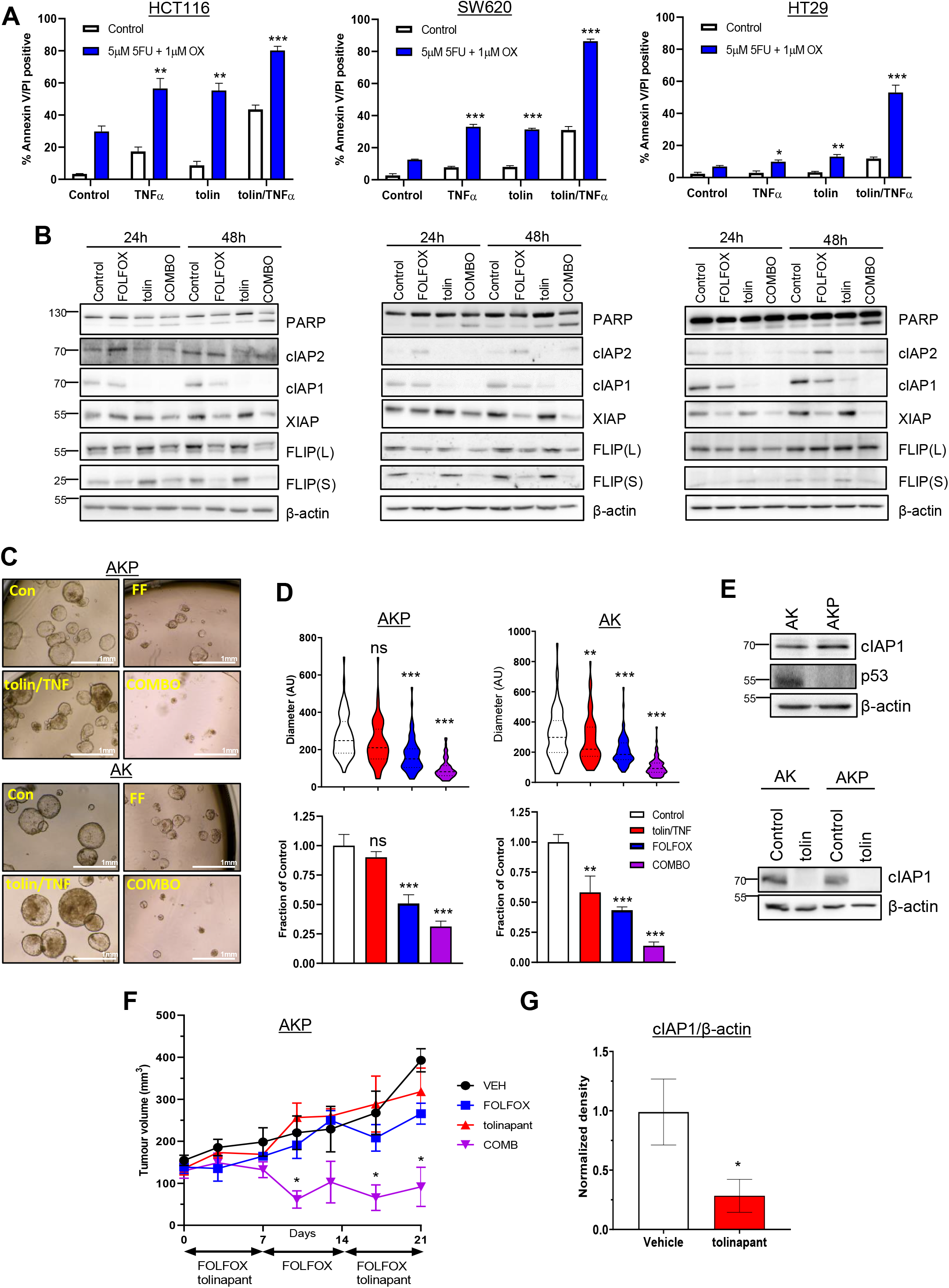
**A)** AnnexinV/PI analysis in HCT116, SW620 and HT29 cells treated with 1ng/ml TNFα, 1μM tolinapant or a combination of tolinapant/TNFα in the presence and absence of 5μM 5FU and 1μM oxaliplatin (OX) for 72h. **B)** Western blot analysis of PARP, cIAP2, cIAP1, XIAP, FLIP(L), FLIP(S) and β-actin in HCT116, SW620 and HT29 cells treated with 5μM 5FU and 1μM oxaliplatin(FOLFOX), 1μM tolinapant and a combination of FOLFOX/tolinapant (COMBO) for 24h and 48h. **C)** Pictures from AKP and AK organoids treated with 1μM tolinapant/1ng/ml TNFα (tolin/TNF), 5μM 5FU and 1μM oxaliplatin (FF) or combination (COMBO) for 72h. Diameter in arbitrary units (AU) from AKP (n=75 for each treatment group) and AK (n=75 for each treatment group) organoids treated with 1μM tolinapant/1ng/ml TNFα (tolin/TNF), 5μM 5FU and 1μM oxaliplatin (FOLFOX) or combination (COMBO) for 72h. **D)** Cell viability assays in AKP and AK organoids treated with 1μM tolinapant/1ng/ml TNFα (tolin/TNF), 5μM 5FU and 1μM oxaliplatin (FOLFOX) or combination (COMBO) for 72h. **E)** Western blot analysis of basal expression of cIAP1, p53 and β-actin in AK and AKP organoids. Western blot analysis of cIAP1 and β-actin in AK and AKP organoids treated with 1μM tolinapant for 24h. **F)** Tumor volume (mm^3^) of AKP organoids transplanted into C57BL/6 mice treated with Vehicle (n=3), 5FU(10mg/kg) + oxaliplatin(1mg/kg) (FOLFOX)(n=4), tolinapant(16mg/kg)(n=4) and combination(n=4). Data shows mean tumor volume per treatment group per timepoint ± sem. **G)** Densitometry from Western blot analysis of cIAP1 normalised to β-actin in AKP organoids transplanted into C57BL/6 mice and treated with Vehicle and 16mg/kg tolinapant. Results were compared using a two-tailed Students t-test, p<0.05 *, p<0.01 ** and p<0.001 ***.

We extended these studies to *Apc*-null, *Kras*-mutant murine CRC organoid models [20-22]. We also examined the impact of p53 status given the high frequency of its mutation in advanced CRC. We observed striking combinatorial effects when both p53-proficient “AK” and p53-deficient “AKP” organoids were co-treated with FOLFOX and tolinapant/TNFα (**Figure 3C/3D**). cIAP1 expression was similar in the 2 organoid models, and tolinapant effectively targeted murine cIAP1 (**Figure 3E**). Taking the AKP model into the *in vivo* setting in immune-competent syngeneic mice, we found that the combination treatment of FOLFOX/tolinapant resulted in significant tumor regression compared to the individual treatments (**Figure 3F**), and *in vivo* on-target activity of tolinapant was confirmed (**Figure 3G**).

### FLIP downregulation in response to FOLFOX sensitises CRC cells to tolinapant

We and others have previously demonstrated the importance of the caspase-8 paralog and regulator *FLIP/CFLAR* as a key mediator of resistance to IAP inhibitors in multiple cancers [23, 24]. Intriguingly, protein expression of the caspase-8 inhibitory short FLIP splice form (FLIP(S)) and XIAP were significantly reduced following FOLFOX treatment in all 5 models (**Figure 3B** and **Figure S3B**). However, as tolinapant is also active against XIAP [25], we hypothesized that the impact of FOLFOX on FLIP was more likely to be responsible for the enhanced effects of tolinapant in FOLFOX-treated CRC cells, particularly in the HCT116 and SW620 *KRAS* mutant models where FLIP downregulation and the extent of synergy were most marked. Moreover, high FLIP mRNA expression corresponded with resistance to tolinapant/TNFα in the extended cell line panel (**Figure 2A/4A**), further supporting the importance of FLIP in modulating response to tolinapant in CRC cells. Somewhat surprisingly, models with high caspase-8 mRNA expression were also more resistant to tolinapant/TNFα, although this can be at least in part explained by the close correlation between caspase-8 and FLIP mRNA expression which are encoded on the same genomic locus (**Figure 4A**). Consistent with a role for FLIP in mediating resistance to tolinapant/TNFα, RNAi-mediated downregulation of FLIP significantly enhanced the induction of apoptosis by tolinapant/TNFα in all but the LoVo model as assessed in short-term cell death (**Figure 4B**) and longer-term cell viability (**Figure 4C**) assays; moreover, these results were confirmed qualitatively by Western blot analysis of PARP cleavage (**Figure 4D/S3C**).

**Figure 4:**
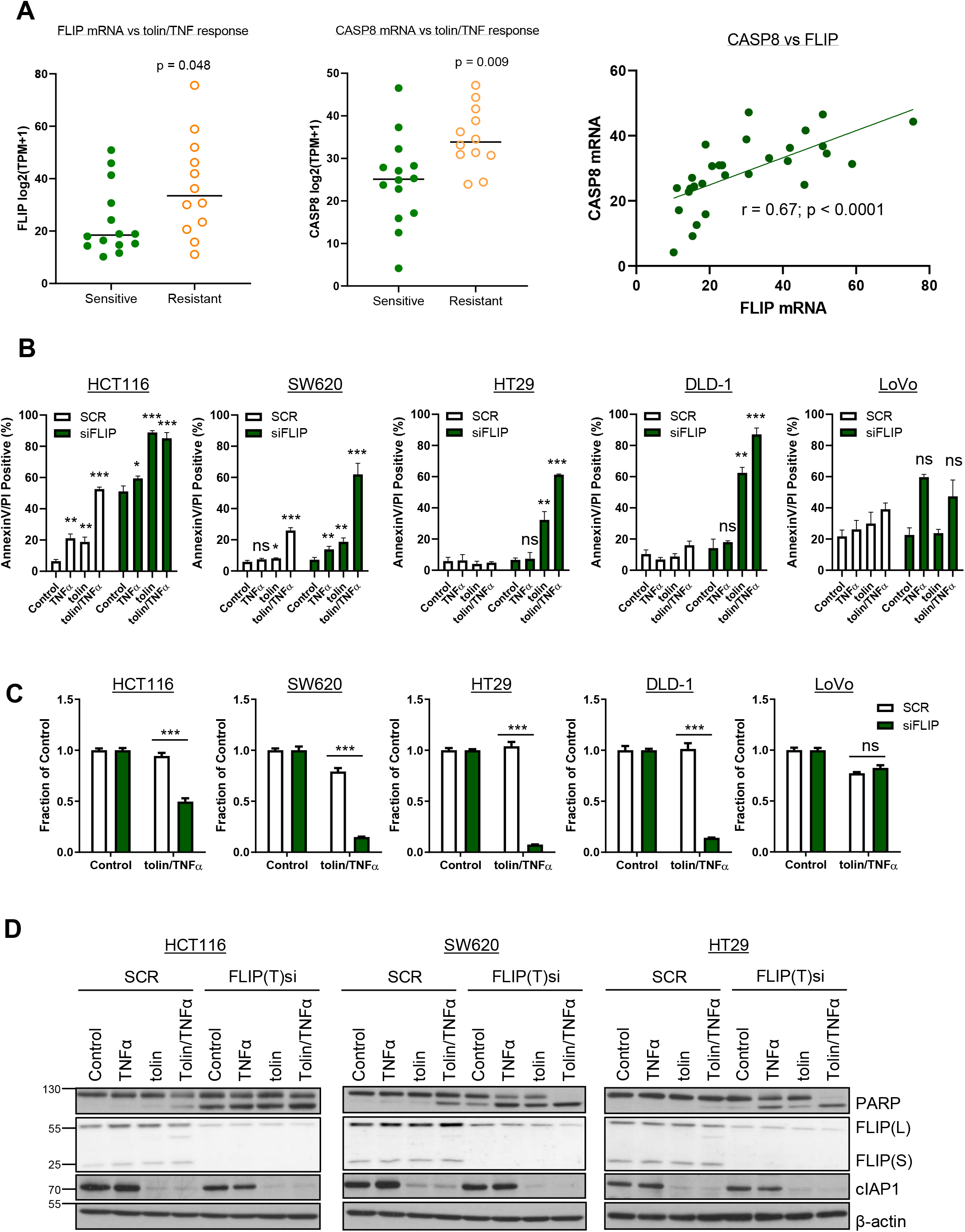
**A)** Expression of Caspase-8 and FLIP mRNA (log2(TPM (transcripts per million) + 1) in CRC cell lines categorised as sensitive or resistant to tolinapant/TNFα in the cell line screen in **Figure 2A**. **B)** AnnexinV/PI analysis in HCT116, SW620, HT29, DLD-1 and LoVo cells transfected with Control (SCR) and FLIP(T) siRNA for 24h prior to treatment with 1μM tolinapant and 1ng/ml TNFα for a further 24h. **C)** Cell viability assays in HCT116, SW620, HT29, DLD-1 and LoVo cells transfected with Control (SCR) and FLIP(T) siRNA for 6h prior to treatment with 1μM tolinapant and 1ng/ml TNFα for a further 24h. **D)** Western blot analysis of PARP, FLIP(L), FLIP(S), cIAP1 and β-actin and in HCT116, SW620 and HT29 cells transfected with Control (SCR) and FLIP(T) siRNA for 24h prior to treatment with 1μM tolinapant and 1ng/ml TNFα for a further 24h. Results were compared using a two-tailed Students t-test, p<0.05 *, p<0.01 ** and p<0.001 ***.

Together these results suggest that FLIP acts as a key mediator of resistance to tolinapant in CRC cells, consistent with downregulation of FLIP being a major contributor to the enhanced efficacy of tolinapant in FOLFOX-treated cells. To formally assess this, we made use of HCT116 cell line models stably expressing retroviral *trans-genes* encoding either FLIP(L), FLIP(S) or a FADD binding deficient mutant form of FLIP(S) (F114A) compromised in its ability to inhibit procaspase-8 homodimerization (**Figure 5A**) [26]. Overexpression of FLIP(S), the more potent caspase-8 inhibitory form of FLIP [27], significantly abrogated apoptosis induced by FOLFOX in combination with tolinapant/TNFα (**Figure 5B**), whereas expression of the FLIP(L) *trans-gene* failed to protect HCT116 cells from apoptosis induced by FOLFOX in combination with tolinapant/TNFα (likely due in part to the relatively small degree of FLIP(L) overexpression in this model; **Figure 5A**). Overexpression of the F114A mutant of FLIP(S) failed to confer protection, indicating that it is FLIP(S)’s interaction with FADD that blocks caspase-8-dependent apoptosis in response to FOLFOX combination with tolinapant/TNFα.

**Figure 5:**
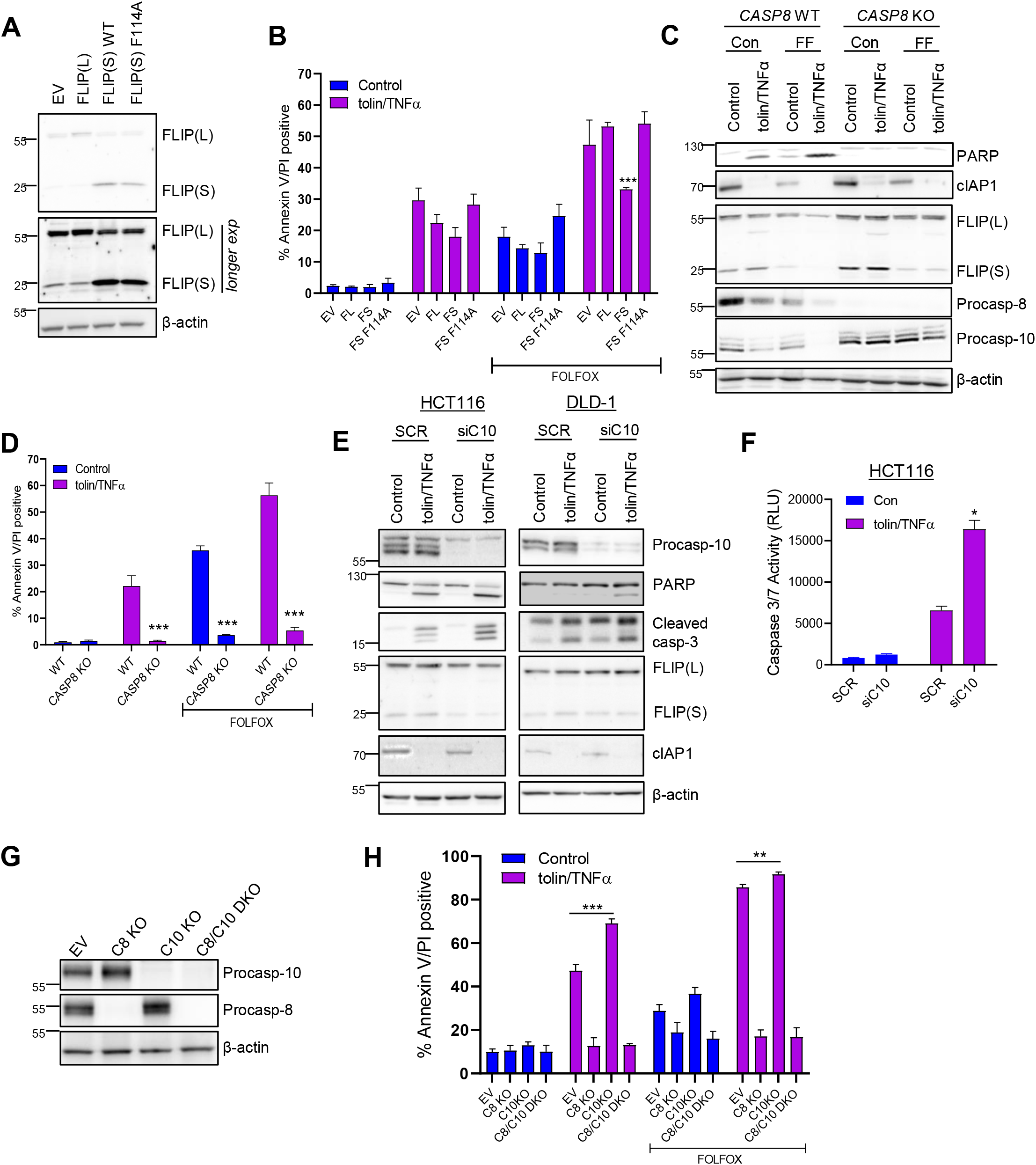
**A)** Western blot analysis of FLIP(L), FLIP(S) and β-actin in HCT116 cells retrovirally overexpressing (o/e) pbabe empty vector (EV), pbabe FLIP(S), pbabe FLIP(S) F114A and pbabe FLIP(L). **B)** AnnexinV/PI analysis of in HCT116 cells retrovirally overexpressing (o/e) pbabe empty vector (EV), pbabe FLIP(S), pbabe FLIP(S) F114A and pbabe FLIP(L) following treatment with 1μM tolinapant/1ng/ml TNFα, 5μM 5FU and 1μM oxaliplatin (FOLFOX) or combination for 48h. **C)** Western blot analysis of PARP, cIAP1, FLIP(L), FLIP(S), Procaspase-8, Procaspase-10 and β-actin in HCT116 WT and HCT116 Caspase 8 KO cells treated with 1μM tolinapant/1ng/ml TNFα (tolin/TNFα), 5μM 5FU and 1μM oxaliplatin (FF) or combination for 48h. **D)** AnnexinV/PI analysis in HCT116 WT and HCT116 Caspase 8 KO cells treated with 1μM tolinapant/1ng/ml TNFα, 5μM 5FU and 1μM oxaliplatin (FOLFOX) or combination for 72h. **E)** Western blot analysis of Procaspase-10, PARP, cleaved caspase-3, FLIP(L), FLIP(S), cIAP1 and β-actin in HCT116 and DLD1 cells transfected with Control (SCR) and caspase-10(siC10) siRNA for 24h prior to treatment with 1μM tolinapant and 1ng/ml TNFα for a further 24h. **F)** Caspase3/7 activity assay in HCT116 cells transfected with Control (SCR) and caspase-10 siRNA (siC10) for 24h prior to treatment with 1μM tolinapant and 1ng/ml TNFα (tolin/TNFα) for a further 24h. **G)** Western blot analysis of Procaspase-10, procaspase-8 and β-actin in HCT116 WT, HCT116 Caspase 8 KO, HCT116 Caspase 10 KO and HCT116 Caspase8/Caspase10 DKO cells. **H)** AnnexinV/PI analysis in HCT116 WT and HCT116 Caspase 8 KO, HCT116 Caspase 10 KO and HCT116 Caspase8/Caspase10 DKO cells treated with 1μM tolinapant/1ng/ml TNFα, 5μM 5FU and 1μM oxaliplatin (FOLFOX) or combination for 48h.

Compared to a HCT116 caspase-8 CRISPR knockout model in which cell death induced by the tolinapant/TNFα-FOLFOX combination was almost completely abrogated (**Figure 5C/D**), significant cell death was still observed in response to the combination in the FLIP(S) overexpressing model (**Figure 5B**), suggesting that additional factors other than FLIP downregulation contribute to the effects of the combination. The role of caspase-8’s other paralog *caspase-10/CASP10* as a regulator of apoptotic cell death is controversial, with both pro- and anti-apoptotic roles described [28-30]. We found that co-treatment of HCT116 cells with tolinapant/TNFα/FOLFOX dramatically reduced the levels of procaspase-10 expression (**Figure 5C**). Loss of the pro-form of procaspases can be indicative of activation and suggested participation of procaspase-10 in the overall mechanism of apoptosis induction. However, in the absence of procaspase-8, relatively little decrease of procaspase-10 was observed (**Figure 5C**). In contrast to the protective effect of loss of procaspase-8, we found that rather than attenuating cell death, siRNA-mediated downregulation of procaspase-10 significantly *enhanced* PARP cleavage and executioner caspase activation in response to tolinapant/TNFα (**Figure 5E/F**). To further investigate this, we generated CRISPR models with either procaspase-8, procaspase-10 or both knocked out (**Figure 5G**). In agreement with the RNAi results, knockout of procaspase-10 enhanced apoptosis induction in response to tolinapant/TNFα in the presence of caspase-8 (**Figure 5H**). These results reveal a hitherto unreported role for procaspase-10 in inhibiting the apoptotic response to IAP antagonists. Furthermore, this is consistent with SW620 cells, which exhibit the lowest levels of procaspase-10 expression, being the most sensitive to tolinapant/TNFα, while DLD-1 cells, with the highest levels of procaspase-10, are highly resistant (**Figure 2B** and **Figure 2F**). Indeed, siRNA-mediated depletion of procaspase-10 also enhanced the sensitivity of DLD-1 cells to tolinapant/TNFα (**Figure 5E**).

### Mechanism of FLIP downregulation in response to FOLFOX

Mechanistically, the effects of FOLFOX on FLIP expression were not due to suppression of transcription; indeed, mRNA expression of FLIP(S) was upregulated at 24h by ~2-fold and FLIP(L) was upregulated by ~5-fold at 48h (**Figure 6A**). Inhibition of histone deacetylases (HDACs) has previously been reported by us and others to result in reduced expression of both splice forms of FLIP, at transcriptional and post-translational levels and augment cell death induced by activators of extrinsic cell death [30-32]. It was therefore notable that FOLFOX treatment downregulated expression of the Class-I HDACs (HDAC-1, −2 and −3) as did tolinapant/TNFα (**Figure 6B**). Moreover, down-regulation of HDAC1 and HDAC3 by both FOLFOX and tolinapant/TNFα was attenuated in *CASP8*-null cells, indicating that these effects are caspase-8-dependent and therefore potentially consequences of apoptosis initiation. In contrast, FOLFOX-mediated downregulation of HDAC2 was caspase-8-independent, suggesting that this is an event upstream of apoptosis initiation.

**Figure 6:**
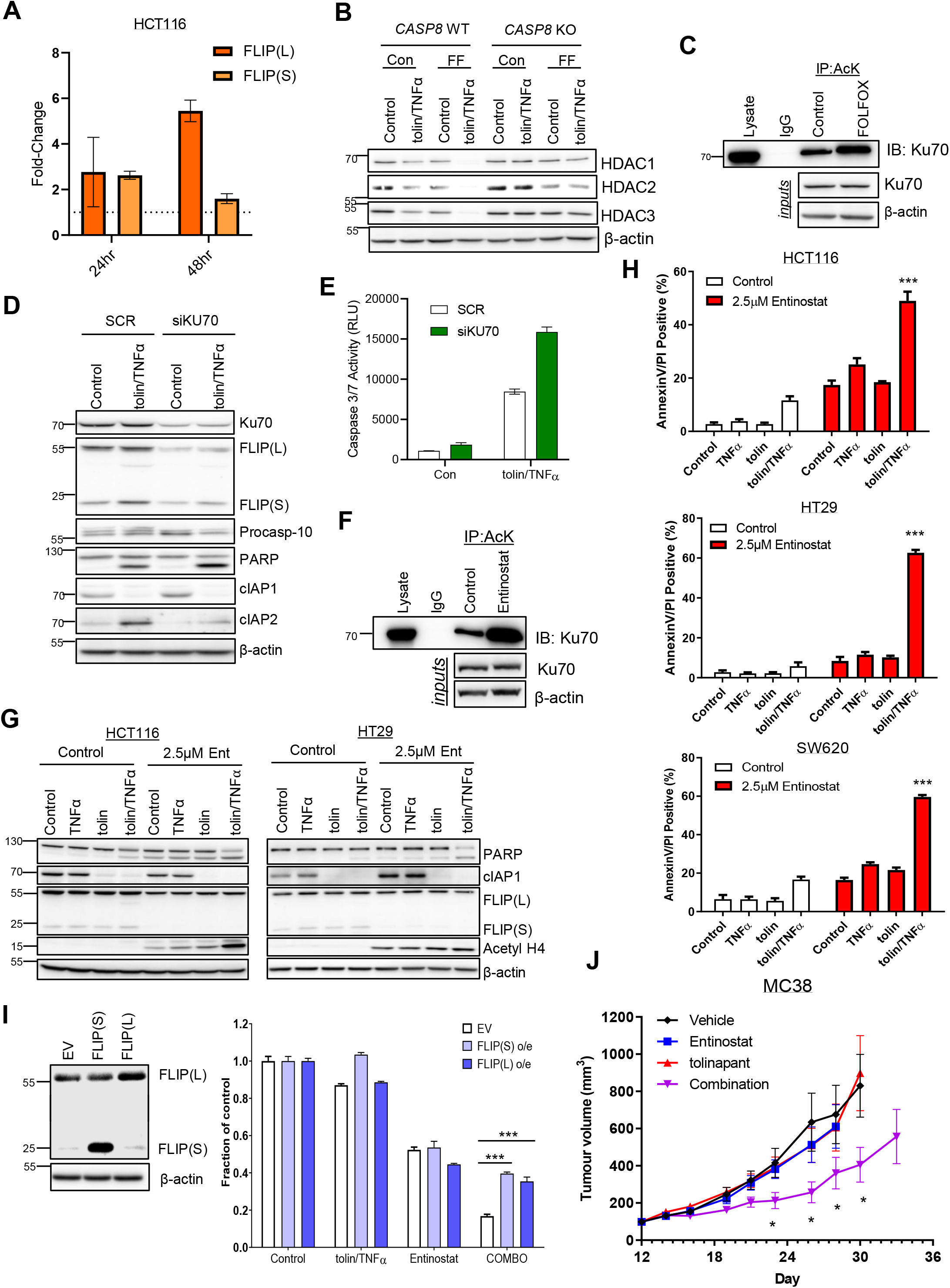
**A)** qPCR analysis of FLIP(L) and FLIP(S) normalised to 18S in HCT116 cells treated with 5μM 5FU and 1μM oxaliplatin for 24h and 48h, relative to untreated control. **B)** Western blot analysis of HDAC1, HDAC2, HDAC3 and β-actin in HCT116 WT and HCT116 Caspase 8 KO cells treated with 1μM tolinapant/1ng/ml TNFα (tolin/TNFα), 5μM 5FU and 1μM oxaliplatin (FF) or combination for 48h. **C)** Western blot analysis of Ku70 following immunoprecipitation of Acetylated Lysine (AcK) from HCT116 cells treated with 5μM 5FU and 1μM oxaliplatin (FOLFOX) for 24h. An IgG isotype control was used as a control. **D)** Western blot analysis of Ku70, FLIP(L), FLIP(S), Procaspase-10, PARP, cIAP1, CIAP2, XIAP and β-actin and in HCT116 cells transfected with Control (SCR) and Ku70 siRNA (siKU70) for 24h prior to treatment with 1μM tolinapant and 1ng/ml TNFα (tolin/TNFα) for a further 24h. **E)** Caspase3/7 activity assay in HCT116 cells transfected with Control (SCR) and Ku70 siRNA (siKU70) for 24h prior to treatment with 1μM tolinapant and 1ng/ml TNFα (tolin/TNFα) for a further 24h. **F)** Western blot analysis of Ku70 following immunoprecipitation of Acetylated Lysine (AcK) from HCT116 cells treated with 2.5μM entinostat for 24h. An IgG isotype control was used as a control. **G)** Western blot analysis of PARP, cIAP1, FLIP(L), FLIP(S), Hyper Acetyl H4 and β-actin in HCT116 and HT29 cells treated with 2.5μM entinostat for 24h then addition of 1μM tolinapant and 1ng/ml TNFα for a further 24h. **H)** AnnexinV/PI analysis in HCT116, HT29 and SW620 cells pretreated with 2.5μM entinostat for 24h then addition of 1μM tolinapant and 1ng/ml TNFα for a further 24h. **I)** Cell viability assay in HCT116 cells retrovirally overexpressing (o/e) pbabe empty vector (EV), pbabe FLIP(S) and pbabe FLIP(L) following pretreatment with 2.5μM entinostat for 24h then addition of 1μM tolinapant and 1ng/ml TNFα for a further 24h. **J)** Tumor volume (mm^3^) in a MC38 syngeneic model treated with Vehicle, entinostat (15mg/kg), tolinapant (16mg/kg) and entinostat/tolinapant combination. Data shows mean tumor volume per treatment group per timepoint ± sem, n=10 for each treatment group.

We previously found that HDAC inhibitors modulate FLIP expression at the post-transcriptional level by inducing acetylation of Ku70, a central component of the non-homologous end-joining (NHEJ) DNA damage repair pathway [32]. In its non-acetylated form, Ku70 binds FLIP and inhibits its proteasomal degradation; however its acetylation in response to HDAC inhibitors blocks this interaction, leading to FLIP degradation. It was therefore notable that FOLFOX caused significant Ku70 acetylation (**Figure 6C**). Moreover, silencing Ku70 not only resulted in decreased levels of FLIP, it also enhanced apoptosis induced by tolinapant/TNFα as assessed by PARP cleavage (**Figure 6D**) and executioner caspase activation (**Figure 6E**).

We also found that entinostat, a Class-I histone deacetylase inhibitor potently induced acetylation of Ku70 (**Figure 6F**) and resulted in downregulation of FLIP(L) and FLIP(S) in all 5 CRC cell lines (**Figure S4A**). Consistent with the effects of FOLFOX and siFLIP, treatment with entinostat resulted in significant sensitisation of HCT116, HT29 and SW620 cells to apoptotic induction by tolinapant/TNFα (**Figure 6G/H**). Similar results were observed in DLD-1 cells; however, LoVo cells remained resistant to tolinapant/TNFα (**Figure S4B/C**), consistent with their lack of sensitization by FLIP silencing (**Figure 4B** and **Figure S3C**). Overexpression of either FLIP(L) or FLIP(S) (**Figure 6I**) abrogated the sensitisation of HCT116 cells to tolinapant/TNFα by entinostat (**Figure 6H**), indicating that entinostat-mediated sensitization is more FLIP-dependent than FOLFOX-mediated sensitization, which was only inhibited by the more highly overexpressed and more potent caspase-8 inhibitor FLIP(S) (**Figure 5B**). Moreover, HCT116 and DLD-1 cells deficient in caspase-8 were resistant to cell death induced by the entinostat/tolinapant/TNFα combination (**Figure S4D**) as was an HCT116 FADD-deficient model (**Figure S4E**).

MC38 murine CRC cancer cells were relatively resistant to tolinapant/TNFα, but were sensitized by FLIP downregulation using siRNA (**Figure S5A/B/C**). Moreover, pre-treatment with entinostat enhanced sensitivity to tolinapant/TNFα *in vitro* (**Figure S5D**), and the combination of entinostat/tolinapant was well tolerated *in vivo* (**Figure S5E**) and resulted in significant tumor retardation compared to either monotherapy (**Figure 6J**). Furthermore, in both the AKP and AK mouse organoid models, entinostat also enhanced the tolinapant/TNFα-induced reduction in organoid size, with the combined effects more marked in the p53-deficient AKP model (**Figure S5F/G**). Overall, these results indicate that Class-I HDAC inhibitors are potential combination partners for tolinapant in CRCs in which FLIP inhibits efficacy.

### Loss or inhibition of caspase-8 leads to necroptosis induction in response to tolinapant/TNFα in RIPK3 positive CRC models

Entinostat was also shown to sensitise HT29 WT cells to tolinapant/TNFα (**Figure 7A**); however, unlike the rescue that was observed in HCT116 and DLD-1 cells (**Figure S4D**), CRISPR-knockout of caspase-8 in HT29 cells resulted in *enhanced* sensitivity to tolinapant/TNFα (and tolinapant alone), even in the absence of entinostat (**Figure 7A**). This effect was abrogated by the necroptosis inhibitor, necrosulfonamide, but not by the pan-caspase inhibitor zVAD, indicating that it is due to the induction of necroptosis rather than apoptosis (**Figure S6A**). Necroptosis is a pro-inflammatory, RIPK1/RIPK3-dependent, but caspase-independent form of programmed cell death. Of the human CRC cell lines profiled for tolinapant activity, only HT29 and SW620 cells expressed RIPK3 at the protein (**Figure 2B**) and mRNA levels (**Figure S6B**). Necroptosis induction was also observed in SW620 *CASP8* KO cells (**Figure S6C**). These results were further supported by Western blot analyses, which demonstrated phosphorylation of RIPK1, RIPK3 and the effector of necroptosis, MLKL (mixed lineage kinase domain-like) in caspase-8-deficient HT29 (**Figure 7B**) and SW620 cells (**Figure S6D**). Moreover, direct assessment of cell death (propidium iodide positivity) further confirmed that the mechanism of cell death induced by tolinapant/TNFα in caspase-8-deificent HT29 (**Figure 7C**) and SW620 cells (**Figure S6E**) was via necroptosis. Furthermore, FADD-deficient HT29 cells were also sensitive to tolinapant/TNFα, a cell death phenotype that was not rescued through inhibition of caspases (**Figure S6F**).

**Figure 7:**
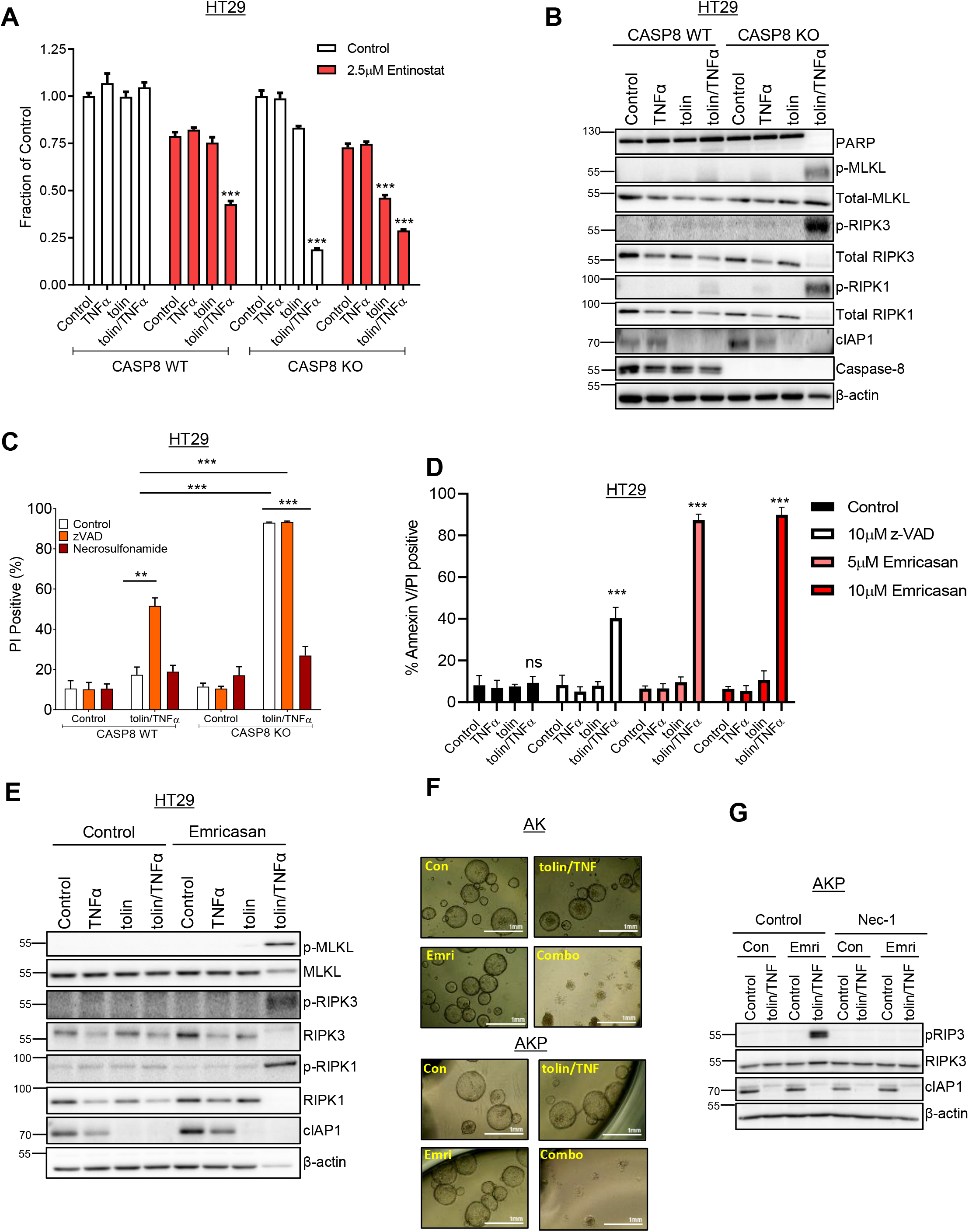
**A)** Cell viability assays in HT29 Caspase 8 WT and HT29 Caspase 8 KO cells pretreated with 2.5μM entinostat for 24h then addition of 1μM tolinapant and 1ng/ml TNFα for a further 24h. **B)** Western blot analysis of PARP, p-MLKL(Ser358), Total MLKL, p-RIPK3(Ser227), Total RIPK3, p-RIPK1(Ser166), Total RIPK1, cIAP1, Caspase 8 and β-actin in HT29 Caspase 8 WT and HT29 Caspase 8 KO cells 24h following treatment with 1μM tolinapant and 1ng/ml TNFα. **C)** Propidium Iodide analysis in HT29 Caspase 8 WT and HT29 Caspase 8 KO cells pretreated for 30 minutes with 10μM z-VAD and 2μM necrosulfonamide then addition of 1μM tolinapant and 1ng/ml TNFα for 24h. **D)** AnnexinV/PI analysis in HT29 cells pretreated with 10μM z-VAD, 5μM emricasan or 10μM emricasan for 30 minutes then addition of 1μM tolinapant and 1ng/ml TNFα for 24h. **E)** Western blot analysis of p-MLKL(Ser358), Total MLKL, p-RIPK3(Ser227), Total RIPK3, p-RIPK1(Ser166), Total RIPK1, cIAP1 and β-actin in HT29 cells pretreated with 10μM emricasan for 30 minutes then addition of 1μM tolinapant and 1ng/ml TNFα for 24h. **F)** Pictures from AK and AKP organoids pretreated with 10μM emricasan for 30 minutes then addition of 1μM tolinapant and 1ng/ml TNFα for 48h. **G)** Western blot analysis of p-RIPK3(Thr231/Ser232), Total RIPK3, cIAP1 and β-actin in AKP organoids pretreated with 10μM Necrostatin-1 and 10μM emricasan (Emri) for 30 minutes before the addition of 1μM tolinapant and 1ng/ml TNFα (tolin/TNF) for 24h. Results were compared using a two-tailed Students t-test, p<0.05 *, p<0.01 ** and p<0.001 ***.

Notably, the pan-caspase inhibitor zVAD induced necroptotic cell death in response to tolinapant/TNFα in parental HT29 cells (**Figure 7C** and **S7A**), but was less efficient at doing so in SW620 cells (**Figure S6C/E**). zVAD also sensitized AK and AKP mouse organoids to tolinapant/TNFα (**Figure S7B**). We therefore investigated the potential of caspase inhibition as a combination partner for tolinapant in RIPK3-positive CRC using emricasan (IDN-6556), a pan-caspase inhibitor already under evaluation in clinical trials in chronic liver disease [33, 34]. Pre-treatment of HT29 or SW620 cells with emricasan resulted in marked sensitisation to necroptosis induced by tolinapant/TNFα (**Figure 7D/E** and **Figure S7C-E**) and was considerably more efficacious than zVAD. These findings were mirrored in the AK and AKP mouse organoid models, where significant combinatorial effects and necroptotic pathway activation were also observed (**Figure 7F/G** and **Figure S7F**).

## Discussion

CRC is the second leading cause of cancer deaths in the Western world, with resistance to programmed cell death contributing to treatment failure [35]. Multiple IAPs are overexpressed in CRC and correlate with therapy resistance and poorer survival rates, making them an attractive target for cancer therapy [36, 37]. Here, we report that stage-III chemotherapy-treated patients with MSS colorectal tumours with elevated expression of cIAP1 or cIAP2 (but not XIAP) have a poor prognosis. This finding makes cIAP1/2-high MSS CRC an attractive subpopulation for IAP-targeted therapeutics.

Pre-clinically, IAP antagonists, including tolinapant, synergise with multiple anti-cancer therapies, including radiotherapy, chemotherapy, TRAIL receptor agonists and hypomethylating agents as well as showing promise in combination with PD-1/PD-L1 blockade [38, 39]. Overall, despite demonstrating potent on-target activity through rapid and sustained downregulation of cIAP1, CRC cell lines showed a limited response to tolinapant alone, although sensitivity was significantly enhanced in many cell lines in the context of a modelled pro-inflammatory microenvironment (co-treatment with TNFα). Moreover, we found that tolinapant/TNFα enhanced cell death induced by FOLFOX, the SoC regimen for CRC, in several human and mouse CRC models *in vitro*, with preclinical anti-tumour activity also demonstrated *in vivo*, suggesting its potential in combination with SoC chemotherapy.

Mechanistically, FOLFOX downregulated expression of FLIP, a pseudo-enzyme paralog of caspase-8 leading to enhanced caspase-8-dependent cell death in response to tolinapant. FLIP negatively regulates apoptosis induction from complex II by inhibiting procaspase-8 homo-dimerization, a prerequisite for the latter’s activation to its apoptosis-inducing catalytic form [17, 40]. The caspase-8/FLIP(L) heterodimer also has catalytic activity, which can inhibit necroptotic signalling from complex IIa/b through the cleavage and inactivation of RIPK1/3 [41]. We and others have previously identified FLIP as a mediator of resistance to IAP antagonist therapy in multiple cancers, including mesothelioma and castrate-resistant prostate cancer [23, 24]. Class-I HDACs (HDAC1/2/3) have been found to be overexpressed in CRC and promote resistance to chemotherapy [42]. Here, we found that FOLFOX treatment resulted in downregulation of the Class-I HDACs and acetylation of their substrate Ku70, an essential component of the non-homologous end-joining DNA damage repair machinery that repairs double-strand breaks, including those induced by DNA damaging chemotherapy, such as FOLFOX [43]. We have previously reported that Ku70 binds to and enhances FLIP stability when in its non-acetylated form [32], a finding corroborated by others [44]. Consistent with this, silencing Ku70 resulted in FLIP downregulation and enhancement of tolinapant/TNFα-induced cell death. Moreover, the oral Class-I HDAC inhibitor, entinostat, which is well-tolerated in humans, also acetylated Ku70 and downregulated FLIP expression resulting in sensitisation to tolinapant in the majority of CRC models. Of note, this combination was well-tolerated *in vivo* and retarded the growth of the MC38 murine CRC model, suggesting potential for Class-I HDAC inhibitors as combination partners for IAP inhibitors in CRC.

Procaspase-10 shares significant homology with procaspase-8: they are 46% identical in their catalytic domains [45], and overexpression of procaspase-10 splice forms 10A and 10D has been shown to re-sensitize caspase-8-deficient cells to TRAIL [28]. However, caspase-10 has also been shown to *inhibit* caspase-8 activation by the Fas/CD95 death receptor, supporting earlier studies in which it was shown that the caspase-8/-10 heterodimer is inactive [29, 46]. Our mechanistic studies are consistent with an inhibitory role for procaspase-10 in regulating apoptosis signalling from the death complexes induced by tolinapant/TNFα. To our knowledge, this is the first study to identify caspase-10 as an inhibitor of Complex-IIa/b.

Necroptosis induced by TNFα signalling is a caspase-independent form of programmed cell death, dependent on RIPK1, RIPK3 and MLKL [47-49]. Necroptosis, unlike apoptosis, results in the activation of inflammatory responses through the release of damage associated molecular patterns (DAMPS) from dying cells, which promote anti-tumor immunity through recruitment of inflammatory cells, maturation of dendritic cells and elicit cross priming of CD8+ T cells [50, 51]. RIPK3 expression is lost in multiple cancers, including many CRCs, and correlates with cancer progression [52-54]. Loss of RIPK3 expression and necroptosis resistance is correlated with *BRAF* gain-of-function mutations, AXL overexpression and RIPK3 promoter hypermethylation [55]. CRISPR-mediated knockout of caspase-8 or FADD in RIPK3-positive CRC models resulted in dramatic induction of necroptosis by tolinapant/TNFα. Thus, if clinical resistance to FOLFOX/tolinapant combination therapy emerges due to procaspase-8 or FADD mutations, BRAF inhibitors, AXL inhibitors or epigenetic modifying agents that upregulate RIPK3 expression may make these resistant tumor cells sensitive to tolinapant-induced necroptosis.

Emricasan is an orally active caspase inhibitor that is well-tolerated in humans [33, 56]. We found that tolinapant and emrciasan co-treatment potently activated necroptosis in the presence of TNFα, but only in RIPK3-positive CRC models. Compared to zVAD, emricasan is a more potent inhibitor of the enzymatic activity of the caspase-8/FLIP(L) heterodimer that cleaves RIPK1 [57]. Our data suggest that tolinapant/emricasan combinations may be effective in pro-inflammatory, RIPK3-positive CRCs.

Multiple IAP antagonists, including tolinapant, are currently in clinical trials, and single agent clinical response was observed predominantly in lymphoma for tolinapant while responses in solid tumours were more limited [12]. Here, we have highlighted the potential of combining tolinapant with FOLFOX in CRC. This is relevant to the majority of patients with CRC as MSS is the predominant disease subgroup in which FOLFOX remains the mainstay of treatment in the neoadjuvant, adjuvant and advanced disease settings. We have also identified FLIP as an intrinsic resistance mechanism to IAP antagonist therapy in CRC, the inhibitory effects of which can be overcome (at least in a subset of disease models) by FOLFOX treatment and the clinically relevant Class-I HDACi, entinostat. Moreover, in RIPK3-positive CRC, the clinically well-tolerated caspase inhibitor, emricasan induces necroptotic cell death. Of note, this may be the preferred mode of cell death in the predominantly immune-cold MSS disease setting due to potent pro-inflammatory effects of necroptosis.

In summary, we have identified three potential combination partners that overcome intrinsic resistance mechanisms to IAP antagonists in definable subsets of CRC, the most immediately actionable of which is of course the combination with SoC chemotherapy, FOLFOX. Such a combination could be particularly beneficial as neoadjuvant treatment of stage-III, cIAP1/2-high MSS disease, which has a particularly poor prognosis when treated adjuvantly with FOLFOX alone.

## Materials and Methods

### Data analysis

MSS stage III treated CRC patient cohort has been extracted from the GSE39582[13] data sets, downloaded from the National Center for Biotechnology Information Gene Expression Omnibus and the expression matrix was processed by collapsing to one probeset per gene using MaxMean from the WGCNA package [58]. The IAP mRNA levels were normally distributed, high and low level of expression was determined using a median split. Kaplan-Meier estimators and Cox proportional hazards regression analysis were assessed using the survival and survminer R packages. Tumour cell type profiling based on gene signature method has been performed by xCELL [59]. The single-sample gene set enrichment analysis (ssGSEA) method was implemented through GenePattern web-tool [60] using the Hallmark gene set molecular signature [61]. Heatmaps were generated after mean z-score normalization. Gene level cell line expression values were assessed from Cancer Cell Line Encyclopaedia (CCLE), curated by the Broad institute [62].

### Compounds

Tolinapant was obtained from Astex Pharmaceuticals (Cambridge, UK). Entinostat, necrosulfonamide and emricasan (IDN-6556) were supplied by Selleckchem (Houston, TX). Z-VAD-fmk was purchased from Calbiochem (Gibbstown, NJ). TNFα was supplied by Prospec (Rehovot, Israel). 5FU and oxaliplatin were obtained from Belfast City Hospital Pharmacy.

### Cell Culture

LoVo, HCT116, DLD-1, SW620 and HT29 cells were purchased as authenticated stocks from ATCC (Teddington, UK). HT29, LoVo and Mc38 cells were maintained in DMEM medium (Invitrogen, Paisley, UK) supplemented with 10% fetal bovine serum (Invitrogen, Paisley, UK). HCT116 and SW620 cells were maintained in McCoys5A medium (Invitrogen, Paisley, UK) supplemented with 10% fetal bovine serum (Invitrogen, Paisley, UK). DLD-1 cells were maintained in RPMI medium (Invitrogen, Paisley, UK) supplemented with 10% fetal bovine serum (Invitrogen, Paisley, UK).

VillinCre^ER^;Apc^fl/fl^;Kras^G12D/+^;Trp53^fl/fl^ (AKP) and VillinCre^ER^;Apc^fl/fl^;Kras^G12D/+^ (AK) organoids were obtained through Dr. Emma Kerr/Prof Owen Sansom (CRUK Career Development Fellowship Ref: A26045). Organoids were cultured in ADF complete media: Advanced DMEM/F12 media (Invitrogen, Paisley, UK), supplemented with 10mM HEPES (Sigma), 2nM L-Glutamine (Invitrogen, Paisley, UK), N2 supplement (Invitrogen, Paisley, UK), B27 supplement (Invitrogen, Paisley, UK), 50ng/ml recombinant human EGF (Peprotech) and 100ng/ml Recombinant murine Noggin (Peprotech). Organoids were harvested in cell recovery solution (Corning, Germany), washed with PBS and organoid pellet resuspended in 1:1 ratio Matrigel (Corning, Germany):Media and cultured in complete media.

### Measuring diameter of organoids

Organoids were cultured and treated as detailed. Pictures of the organoids were taken with an EVOS XL core cell imaging system (Thermo Fisher Scientific) with x10 magnification. Diameter of organoids, in arbitrary units (AU), were determined utilising ImageJ software.

### Cell line screen

36-cell line screen was carried out by Chempartner (China). Each cell line was tested once in duplicate following incubation with 10 different concentration of tolinapant (0.0005 – 10μM) alone and in combination with TNFα (1ng/ml) for 72h. Cell viability was assessed by CellTitreGlo.

### Western Blotting

Western Blotting was carried out as described previously [23]. Mcl-1, XIAP, BAX, SMAC, RIPK1, RIPK3, MLKL, Acetylated Lysine, pMLKL(Ser358), pRIPK3(Ser227), pRIPK1(Ser166) and PARP antibodies were purchased from Cell Signaling Technology (Danvers, MA). cIAP1, cIAP2 and Caspase-8 antibodies were purchased from Enzo (Exeter, UK). FLIP antibody was from AdipoGen (San Diego, CA) and Hyperacetylated H4 was from Millipore (Gillingham, UK). FADD antibody was purchased from BD Biosciences, Caspase-10 from MBL (Woburn, MA), Ku70 from Santa Cruz (Dallas, Texas) and β-actin from Sigma.

### siRNA transfections

ON-TARGET plus SMART pool Murine cFLIP siRNA and SCR, cFLIP(L), cFLIP(S), cFLIP(T) and caspase-10 siRNA’s was purchased from Dharmacon (Chicago, IL). siRNA transfections were carried out with Lipofectamine RNAiMAX (Invitrogen, Paisley, UK) transfection reagent according to manufacturer’s instructions. siRNA sequences: SCR: UUCUCCGAACGUGUCACGU, FLIP(L): AACAGGAACTGCCTCTACTT, FLIP(S): AAGGAACAGCTTGGCGCTCAA, FLIP(T): AAGCAGTCTGTTCAAGGAGCA. Ku70 siRNA transfections were carried out as previously described [32].

### Quantitative PCR (qPCR)

RNA was extracted using High Pure RNA Isolation kit (Roche, Burgess Hill, UK) according to manufacturer’s instructions. cDNA was synthesised using Transcriptor First Strand cDNA synthesis kit (Roche, Burgess Hill, UK) according to manufacturer’s instructions. qPCR was carried out using Syber green on LC480 light cycler (Roche, Burgess Hill, UK) according to manufacturer’s instructions. Primer sequences: cFLIP(L): F:CCTAGGAATCTGCCTGATAATCGA, R:TGGGATATACCATGCATACTGAGATG, FLIP(S): F:ATTTCCAAGAATTTTCAGATCAGGA, R:GCAGCAATCCAAAAGAGTCTCA, 18S: F:GTAACCCGTTGAACCCCATT, R:CCATCCAATCGGTAGTAGCG, RIPK3: GCCTCCACAGCCAGTGAC, TCGGTTGGCAACTCAACTT, Murine cFLIP: F:GCAGAAGCTCTCCCAGCA, R:TTTGTCCATGAGTTCAACGTG, Murine 18S: F:GCAATTATTCCCCATGAACG, R:GGGACTTAATCAACGCAAGC.

### Cell Line Generation

HCT116 cells overexpressing cFLIP(S) or cFLIP(L) were generated as previously described [23]. HCT116, DLD1, LoVo, HT29 and SW620 caspase-8 CRISPR cell lines were generated by retroviral infection with pLentiCRISPR with guideRNA AAGTGAGCAGATCAGAATTG, a kind gift from Prof. Galit Lahav [63]. HCT116 and HT29 FADD CRISPR cell lines were generated by retroviral infection with pLentiCRISPR with guideRNA GCGGCGCGTCGACGACTTCG. Mixed population cell lines were established following selection with 1μg/ml puromycin. Cells were clonally selected for complete knockdown by Western blot analysis. HCT116 Caspase10 CRISPR cell lines were generated as previously described [30].

### Immunoprecipitations

Caspase-8 and Acetylated Lysine immunoprecipitations were carried out as previously described [23, 32].

### Cell surface flow cytometry

TNFR1 cell surface staining was carried out a previously described [23].

### Cell viability assay

2D cell viability was assessed using CellTiter-Glo Luminescent Assay (Promega, Madison, WI) according to manufacturer’s instructions. 3D cell viability was determined with CellTitre-Glo 3D Cell Viability Assay (Promega, Madison, WI), according to manufacturer’s instructions.

### AnnexinV/PI Staining

FITC-tagged AnnexinV (BD-Biosciences), PI(Sigma) and Hoesch stain (Invitrogen) were used to assess cell death. AnnexinV/PI staining was assessed by high content microscopy on an Array Sacn XTI microscope (Thermo Scientific).

### In vivo experiments

2×10^6^ HCT116, HT29 and DLD-1 cells in PBS:Matrigel(Corning, Germany) suspension were implanted single flank into 8-10 week old male BALB/c nude mice. When tumors of approx.100mm^3^ were established, mice were randomized and treated with vehicle and tolinapant. Tolinapant was administered orally with H_2_O as a vehicle daily, one week on, one week off, one week on.

Mc38 syngeneic experiment was carried out by Crown Bioscience (Loughborough, UK). Mc38 cells were injected subcutaneously injected into C57BL-6 mice (single flank). When tumors of approx.100mm^3^ were established, mice were randomized and treated orally with vehicle, entinostat(15mg/kg), tolinapant(16mg/kg) or combination of entinostat and tolinapant. The vehicle for entinostat was 30% cyclodextrin and for tolinapant was H_2_O. Mice were treated for a total of three weeks with entinostat administered twice a week and tolinapant daily on a one week on, one week off, one week on schedule.

50 AKP organoids in PBS:Matrigel(Corning, Germany) suspension were subcutaneously injected into C57BL-6J mice and treatment stated when average tumor size ~150mm^3^. Mice were treated with 10mg/kg 5FU (vehicle:PBS) and 1mg/kg oxaliplatin (vehicle:H_2_O) intraperitoneally and 16mg/kg tolinapant (vehicle:H_2_O) orally. Mice were treated for three weeks: once a week with oxaliplatin, daily with 5FU and tolinapant daily with a one week on, one week off, one week on schedule.

All procedures were carried out in accordance with the Animals (Scientific Procedures) Act, 1986 under project licences PPL 2590b and PPL 2874 and maintained as previously described [24]. Tumor volume was assessed using digital calipers, three times a week. Tumor volume was calculated by the following formula: shortest tumor diameter^2^ × longest tumor diameter × 0.5.

## Statistical Analysis

Results were compared using a two-tailed Students t-test, p<0.05 *, p<0.01 ** and p<0.001 ***.

## Acknowledgements

This work was supported by a studentship (KS) sponsored by The Northern Ireland Department for the Economy (NI DfE), and by funding from a CRUK Program grant (DBL/SMcD) C11884/A24367, a CRUK Experimental Cancer Medicine Centre (ECMC) award (CL/VC/DL) C36697/A25176, and a CRUK Career Development Fellowship (EK/WMcD) C61288/A26045. Additional support (DBL, CMcC) was obtained from NI DfE (SFI-DEL 14/1A/2582). TA and ML were supported by an HDR-UK grant (JHR1157-100/1230). DBL was also in receipt of direct funding from Astex Pharmaceuticals.

## Supplementary Figure Legends

**Supplementary Figure 1:** Additional analysis of MSS, Stage III patients who received adjuvant fluorouracil-based treatment extracted from GSE39582 cohort. **A)** Distribution of BIRC2, BIRC3 and XIAP gene expression across the cohort. Red lines show the average, while the blue lines denotes the median value that have been used to determine High and Low category for each gene. **B)** Kaplan-Meier plots of 5-year overall survival comparing the difference in survival probability based on XIAP expression levels. **C)** Distribution of BRAF, KRAS, p53 mutations (wild type = WT and mutant=MT) in BIRC2 and BIRC3 High or Low patients. **D)** Difference of BIRC2, BIRC3 and XIAP expression values in KRAS mutant (M) and wild type (WT) patients. **E)** Effect of KRAS mutation in patients. Kaplan-Meier plots of 5-year overall is shown with the log-rank test, the hazard ratio (HR) with its 95% confidential interval and the Wald test of significance.

**Supplementary Figure 2: A)** Mutational and microsatellite instability status of DLD-1 SW620, LoVo, HT29 and HCT116 CRC cell lines. wt (wild type), MSI (microsatellite instability), MSS (microsatellite stable). Adapted from Ahmed, D., et al (2013)[64]. **B)** Western blot analysis of cIAP1, cIAP2, XIAP and β-actin in LoVo and SW620 cells following treatment with 1μM, 100nM, 10nM, 1nM and 0.1nM tolinapant for 24h. **C)** Western blot analysis of c-IAP1, cIAP2, XIAP, FLIP(L), FLIP(S), procaspase-8, RIPK1 and β-actin in HCT116 and LoVo cells at 1h, 3h, 6h,10h and 24h following treatment with 1μM tolinapant. **D)** Western blot analysis of RIPK1 and procaspase-8 following immunoprecipitation of caspase-8 from LoVo and SW620 cells treated with 1ng/ml TNFα, 1μM tolinapant and a combination of TNFα/tolinapant for 3h in the presence of 20μM z-VAD-fmk. An IgG isotype control was used as a control.

**Supplementary Figure 3: A)** AnnexinV/PI analysis in DLD-1 and LoVo cells treated with 1ng/ml TNFα, 1μM tolinapant or a combination of TNFα/tolinapant in combination with 5μM 5FU and 1μM oxaliplatin (OX) for 72h. **B)** Western blot analysis of PARP, cIAP2, cIAP1, XIAP, FLIP(L), FLIP(S) and β-actin in DLD-1 and LoVo cells treated with 5μM 5FU and 1μM oxaliplatin(FOLFOX), 1μM tolinapant and a combination of FOLFOX/tolinapant (COMBO) for 24h and 48h. **C)** Western blot analysis of PARP, FLIP(L), FLIP(S), cIAP1 and β-actin and in DLD-1 and LoVo cells transfected with Control (SCR) and FLIP(T) siRNA for 24h prior to treatment with 1μM tolinapant and 1ng/ml TNFα for a further 24h. Results were compared using a two-tailed Students t-test, p<0.05 *, p<0.01 ** and p<0.001 ***.

**Supplementary Figure 4: A)** Western blot analysis of FLIP(L), FLIP(S), cIAP1, Hyper Acetyl H4 and β-actin in HCT116 HT29, SW620, DLD-1 and LoVo cells treated with 0.5μM, 1μM, 2.5μM, 5μM entinostat for 24h. **B)** AnnexinV/PI analysis in DLD-1 and LoVo cells pretreated with 2.5μM entinostat for 24h then addition of 1μM tolinapant and 1ng/ml TNFα for a further 24h **C)** Western blot analysis of PARP, cIAP1, FLIP(L), FLIP(S), Hyperacetylated Histone 4 (H4) and β-actin in SW620, DLD-1 and LoVo cells treated cells pretreated with 2.5μM entinostat for 24h then addition of 1μM tolinapant and 1ng/ml TNFα for a further 24h. **D)** Western blot analysis of PARP, Caspase-8, FLIP(L), FLIP(S), cIAP1, Acetyl Histone 4(H4) and β-actin in HCT116 and DLD-1 WT and Caspase-8 KO cells pretreated with 2.5μM entinostat(Ent) for 24h then addition of 1μM tolinapant and 1ng/ml TNFα for a further 24h. **E)** AnnexinV/PI assay in HCT116 WT and HCT116 FADD KO cells pretreated with 2.5μM entinostat for 24h then addition of 1μM tolinapant (tolin) and 1ng/ml TNFα (TNF) for a further 24h. Results were compared using a two-tailed Students t-test, p<0.05 *, p<0.01 ** and p<0.001 ***.

**Supplementary Figure 5: A)** Western blot analysis of cIAP1 and β-actin in MC38 cells treated with 1μM, 100nM, 10nM, 1nM and 0.1nM tolinapant for 24h. **B)** Cell viability assay in MC38 cells 72h following treatment with 1μM tolinapant, 1ng/ml TNFα and a combination of tolinapant/TNF. **C)** Cell viability assay in MC38 cells transfected with scrambled control (SCR) or FLIP (mFLIPsi) siRNA for 24h followed by treatment with 1μM tolinapant and 1ng/ml TNFα for a further 24h. qPCR analysis for FLIP normalised to 18S in Mc38 cells transfected with scrambled control (SCR) and FLIP(mFLIPsi) siRNA for 24h. **D)** Cell viability assays in MC38 cells pretreated with 2.5μM entinostat for 24h then addition of 1μM tolinapant and 1ng/ml TNFα or a combination of tolinapant/TNF for a further 48h. **E)** Analysis of body weight in MC38 syngeneic model illustrated as % of weight relative to initiation of treatment. **F)** Pictures of AKP and AK organoids pretreated with 2.5μM entinostat(Ent) followed by addition of 1μM tolinapant and 1ng/ml TNFα(tolin/TNF) for a further 48h. **G)** Diameter in arbitrary units (AU) of AKP (n=175 for each treatment group) and AK (n=50 for each treatment group) organoids pretreated with 2.5μM entinostat(Ent) followed by addition of 1μM tolinapant and 1ng/ml TNFα(tolin/TNF) for a further 48h. Results were compared using a two-tailed Students t-test, p<0.05 *, p<0.01 ** and p<0.001 ***.

**Supplementary Figure 6: A)** Cell viability assays in HT29 WT and HT29 Caspase 8 CRISPR cells pretreated for 30 minutes with 10μM z-VAD and 2μM necrosulfonamide then addition of 1μM tolinapant and 1ng/ml TNFα for 24h. **B)** qPCR analysis of RIPK3 normalised to 18S in HT29, SW620, DLD-1, HCT116 and LoVo cells. **C)** Cell viability assay in SW620 WT and SW620 Caspase 8 CRISPR cells pretreated for 30 minutes with 10μM z-VAD and 2μM necrosulfonamide then addition of 1μM tolinapant and 1ng/ml TNFα for 48h. **D)** Western blot analysis of PARP, p-MLKL(Ser358), Total MLKL, p-RIPK1(Ser166), Total RIPK1, cIAP1, Caspase 8 and β-actin in SW620 WT and SW620 Caspase-8 KO cells treated with 1μM tolinapant and 1ng/ml TNFα for 24h. **E)** Propidium Iodide analysis in SW620 Caspase 8 WT and SW620 Caspase 8 KO cells pretreated for 30 minutes with 10μM z-VAD and 2μM necrosulfonamide then addition of 1μM tolinapant and 1ng/ml TNFα for 24h. **F)** Cell viability assays in HT29 WT and HT29 FADD KO cells pretreated for 30 minutes with 10μM z-VAD then addition of 1μM tolinapant and 1ng/ml TNFα for 24h. Results were compared using a two-tailed Students t-test, p<0.05 *, p<0.01 ** and p<0.001 ***.

**Supplementary Figure 7: A)** Western blot analysis of p-MLKL(Ser358), Total MLKL, p-RIPK3(Ser227), Total RIPK3, p-RIPK1(Ser166), Total RIPK1, cIAP1 and β-actin in HT29 cells pretreated for 30 minutes with 10μM z-VAD then addition of 1μM tolinapant and 1ng/ml TNFα for 24h. **B)** Pictures from AK (n=100 for each treatment group) and AKP (n=200 for each treatment group) organoids pretreated with 10μM z-VAD for 30 minutes followed by tolinapant/1ng/ml TNFα (tolin/TNF) for 72h. Diameter in arbitrary units (AU) from AKP and AK organoids pretreated with 10μM z-VAD for 30 minutes followed by tolinapant/1ng/ml TNFα (tolin/TNF) for 72h. **C)** Western blot analysis of p-MLKL(Ser358), Total MLKL, Total RIPK3, p-RIPK1(Ser166), Total RIPK1, cIAP1 and β-actin in SW620 cells pretreated with 10μM emricasan for 30 minutes then addition of 1μM tolinapant and 1ng/ml TNFα for 24h. **D)** AnnexinV/PI staining of SW620 cells pretreated with 10μM z-VAD, 5μM emricasan or 10μM emricasan for 30 minutes then addition of 1μM tolinapant and 1ng/ml TNFα for 24h.**E)** Cell viability assays in HT29 and SW620 cells pretreated with 10μM z-VAD, 5μM emricasan or 10μM emricasan for 30 minutes then addition of 1μM tolinapant and 1ng/ml TNFα for 24h. **F)** Cell viability assays in AK and AKP organoids pretreated with 10μM emricasan for 30 minutes then addition of 1μM tolinapant and 1ng/ml TNFα for 48h. Results were compared using a two-tailed Students t-test, p<0.05 *, p<0.01 ** and p<0.001 ***.

